# LoFi drafts to map to: 4 haplotype-resolved Cannabis genomes enable characterization of large structural variants

**DOI:** 10.64898/2026.01.19.700373

**Authors:** Brett Pike, Anders Gonçalves da Silva, Wilson Terán

## Abstract

We present fully-phased, chromosome-scale genome assemblies of 4 genotypes of *Cannabis sativa.* These assemblies were built from Oxford Nanopore R9.4.1 long reads, which previously have been considered insufficiently accurate for proper phasing. Contigs produced by the Phased Error Correction and Assembly Tool (PECAT), in combination with Hi-C libraries, were used by GreenHill to develop intermediate data structures that permit accurate phasing of the dual contigs, which were then scaffolded by the advanced algorithm of Yet another Hi-C Scaffolder (YaHS). These assemblies, while low in QV, are comparable to recent HiFi assemblies in their contiguity and gene content, and also show good macrosynteny with them. We compare these 8 haplotypes with 77 others recently produced and present a phylogenetic analysis, as well as a first draft of the Cannabis pan-NLRome.

**Core:** We assembled four fully-phased and chromosome-scale diploid genomes of Cannabis sativa, using Oxford Nanopore Technology readsets. These new assemblies are comparable to recent PacBio HiFi assemblies in terms of contiguity and gene content. We present a phylogenomic analysis, using whole-genome alignments after including 77 other publicly available Cannabis genomes, as well as a draft pan-NLRome.

**Gene and Accession Numbers:** Assemblies are archived at NCBI as BioProjects PRJNA1301983 (ANC), PRJNA1301963 (HAW), PRJNA1301984 (SRI), and PRJNA1301985 (TRC).

Assemblies, annotations, and Supplemental Tables are also available on Zenodo: https://doi.org/10.5281/zenodo.16456638.

## Introduction

In the era of its legalization, genetic knowledge of Cannabis is rapidly approaching a level comparable to that of other minor crops. In the last decade, more than 30,000 papers have been published on Cannabis, representing more than two-thirds of total output on the subject (NORML 2023). More recently, several chromosome-scale assembled genomes (Braich *et al*. 2020; Gao *et al*. 2020; Grassa *et al*. 2021; Phylos Bioscience, Inc. 2022; Lim 2023), as well as 181 PacBio assemblies and two pangenome graphs (Lynch *et al*. 2025), have recently been released into the public domain.

With the advent of longer and more accurate sequencing reads, brought by Third Generation Sequencing platforms, it is clear that within the same species, plant genomes harbor tremendous structural diversity. In both corn (Goel *et al*. 2019) and cannabis (Pike, Kozik and Terán 2025), aligning two good-quality assemblies will reveal about one third of the genome to be highly diverged. Therefore, in order to accurately characterize large SVs, it is preferred to assemble a genome *de novo*, without incorporating positional or sequence information from another draft of moderate similarity.

Here, we present haplotype-resolved, chromosome-scale genome assemblies for four genotypes of Cannabis, for a total of 8 new assemblies. The use of ultra-long Oxford Nanopore Technology (ONT) libraries allows these assemblies to span repeats that may not be resolved with reads whose lengths are limited by PCR processivity. Subsequent scaffolding using Hi-C libraries both phased contigs and also raised them close to chromosome scale. These new assemblies are highly contiguous, which facilitates the discovery of large structural variants.

Previously we characterized the Nucleotide-binding, Leucine-rich Repeat (NLR) family of pathogen resistance genes, as well as the terpene synthase gene family complements of both haplotypes of a wide cross. Here, we extend that gene family analysis to include these new 8 haplotypes, together with 77 other high-quality genome assemblies, which enables us to visualize the population structure, characterize genetic admixture, and present here the first draft of the Cannabis NLRome.

## Materials and Methods

Example scripts for assembly and phylogenetic analyses may be found at https://github.com/COMInterop/lighthouse.

### Germplasm Selection

Four Cannabis genotypes were selected for *de novo* assembly: Anders CBD (ANC), Hawthorne OG (HAW), Sri Lankan (SRI), and Travis CBD (TRC). Hawthorne OG and Sri Lankan are marijuana (MJ) types, whose primary cannabinoid is psychoactive tetrahydrocannabinol (THC), and Anders CBD and Travis CBD are high-cannabinoid hemp (HC_HEMP) types, whose primary cannabinoid is non-psychoactive cannabidiol (CBD). Hawthorne OG, a variant of OG Kush, is a popular variety in adult-use cannabis dispensaries in California and British Columbia, due to its high THC content. The Sri Lankan was chosen as a rare equatorial Asian variety, which may represent a distinct and unusual lineage. Anders CBD and Travis CBD are grown for medicinal CBD production and have very low THC content. For convenience, we refer to these as the Lighthouse genotypes. Our comparative analysis denotes the two haplotypes as ‘a’ and ‘b’.

For comparison, we include the Punto Rojo (PR) and Cherry Pie (CP) haplotypes assembled by us previously, i.e. the Medcann genotypes, as well as 75 assemblies downloaded from the website of the Salk Institute Cannabis Pangenome, selected to represent the diversity of fibre hemp (HEMP), HC_HEMP, and ‘sativa’ and ‘indica’ MJ. 46 are phased haplotypes built from PacBio HiFi, and the others are pseudohaploid: 24 PacBio CCS, 4 PacBio CLR, and 1 Oxford Nanopore. The Salk genotypes are summarized in Supplemental Table 1, where codes not appended with ‘a’ or ‘b’ are pseudohaploid.

### DNA purification

Samples were delivered to the National Research Council in Toronto, Canada. HMW DNA was purified from clean nuclei via a salting-out method (Ramsay *et al*. 2021) and then size-selected above 50kb with a Blue Pippin (Sage Science, Beverly, Massachusetts, USA). As well, Illumina libraries were prepared using the Illumina PCR-free kit for polishing, and the Arima v1.0 kit for Hi-C.

### Sequencing

Each sample was sequenced in two cells on the Oxford Nanopore Technology PromethION platform, except for Sri Lankan, which was sequenced on three MinION cells. During method optimization, Hawthorne OG was also sequenced on 2 MinIONs. All employed the R9.4.1 pore and were base-called with Guppy 6 in HAC mode. The 8 Illumina libraries were sequenced in paired-end mode in one NovaSeq lane, with read lengths of 150 and 156 bp for polishing and Hi-C, respectively.

### Assembly

ONT reads were filtered for adapters with Porechop (Wick 2018). Due to the scant data for Sri Lankan, the failed reads (<Q8) were included for assembly as we found previously that error-correction was able to rescue usable portions from them (Pike, Kozik and Terán 2025). The Illumina readsets were filtered with BBDuk (Bushnell 2018) to remove adapters and reads below quality 14 or length 21.

Readsets were assembled into contigs with PECAT (Nie *et al*. 2024). Relative to defaults, correction input was raised from 80x to 112x, and assembly input from 80x to 88x. The block size for correction and assembly was raised from 4Gb to 400Gb to permit true all-vs-all alignment. Among parameters passed to minimap2 (Li 2018), the minimizer size was raised from 15 to 19 for error correction (-k19). For the correction and assembly phases, we used ‘-f 0.005’ and ‘-f 0.001’ to discard the 0.5% and 0.1% most abundant kmers. Both phases used ‘-I 400g -K 400g’ so as to rely on one-part indexes and a single minibatch. For assembly, the -r parameter was raised from 500bp to 20kb. For PECAT phasing, ‘phase_method=1’ was set to use only Clair3 (Zheng *et al*. 2022) and not the FSA method, and corrected reads were mapped with ‘-x map-ont -c --secondary=no --rmq=yes -r 1000,250k -K400g −2’. Contig output was filtered above 4 Kb.

Diploid-aware long-read polishing took place within PECAT using Racon (Vaser *et al*. 2017) with default settings. Next, the PECAT process for mapping uncorrected haplotypic reads occurred, but variant calling was performed with Clair3 instead of Medaka, with option ‘--haploid precise’ to only retain 1/1 variants. All Clair3 calls, including those marked as ‘LowQual,’ were used to generate a consensus with bcftools.

Following assembly, the primary and alternate drafts were each cleaned of duplicates with purge_haplotigs (Roach, Schmidt and Borneman 2018) and then concatenated. In the case of Sri Lankan, which had much less coverage, the haplotigs purged from the primary were concatenated with the alternate draft, which was then haplotig-purged prior to concatenating with the primary.

Illumina reads were mapped to the concatenated primary + alternate (pri-alt) assembly with BWA-MEM2 (Vasimuddin *et al*. 2019) with default settings. Polishing was as described previously (Pike, Kozik and Terán 2025). We called 1/1 variants with Clair3 and repeated the procedure. 40-mers and 26-mers were counted in the Illumina reads with ntHits (Warren *et al*. 2019) with options ‘-b 36 --outbloom --solid’. The contigs were then polished with ntEdit (Warren *et al*. 2019). Four rounds of polishing transpired with kmers of size 40, 26, 40, and 26.

The intermediate pseudohaploid contigs were mapped to ‘Sour Diesel B’ (SODLb) (Lynch *et al*. 2025), which we use as a reference, and binned per chromosome, based on each contig’s longest alignment with the reference. Each chromosome was scaffolded separately with HapHiC (Zhou, McCarthy and Durbin 2023), based on information derived from mapping the Hi-C reads to the complete draft with BWA-MEM2, so that the same BAM was used for all 10 HapHiC events. The final diploid contigs, as well as ONT and Hi-C reads, were then mapped to the pseudohaploid chromosomes and binned similarly. The Hi-C readset was filtered to remove pairs where both reads did not map to contigs assigned to the same chromosome. Then, GreenHill ((Ouchi, Kajitani and Itoh 2023) was used to produce preliminary scaffolds for each chromosome, with ONT and filtered Hi-C reads provided as input.

The per-chromosome Greenhill scaffolds were next used to separate the diploid contigs between haplotypes 0 and 1. Each putative haplotype contig set was first mapped to its GreenHill scaffolds with minimap2. Following visualization as a dotplot, ‘scrap’ contigs were discarded and, when two allelic contigs mapped to one haplotype, one was reassigned to the other. For each chromosome, each haplotype’s contigs were scaffolded with YaHS (Zhou, McCarthy and Durbin 2023), again using one whole-genome Hi-C mapping for all 20 YaHS events. After scaffolds were error-corrected within YaHS, and in rare cases broken to resolve spurious telomere:telomere joins, they were aligned to SODLb, reverse-complemented where necessary, and joined manually, with 200 Ns between scaffolds, so as to distinguish from the 100 Ns between contigs within a scaffold. Finally, pseudochromosomes were aligned against the genome of cs10 (Grassa *et al*. 2021), the initial NCBI reference for Cannabis, and where necessary reverse-complemented to match its orientation. For a small number of chromosomes, where the dotplot showed large gaps in the second haplotype, presumably due to homozygosity, some contigs were included in both haplotypes.

### Annotation

Annotation for all genes, resistance genes, and terpene synthases was as described previously (Pike, Kozik and Terán 2025). Briefly, annotations from cs10 were applied with Liftoff (Shumate and Salzberg 2021) with exonic stringency of 99%. Terpene synthase genes (TPS) were separately annotated with Liftoff, at exonic stringency of 50%, in order to find any reasonable paralogs. Resistance genes belonging to the Nucleotide-binding Leucine-rich Repeat (NLR) family were annotated by taking the intersection of two prediction pipelines: the NLR-Annotator (Steuernagel *et al*. 2020), and a custom HMM profile built by aligning the nucleotide binding sites (NBS) of NLRs found in the cs10 reference.

Repeat content was characterized with panEDTA (Ou *et al*. 2024).

### Alignment

Pairwise alignments were performed with minimap2 with options ‘-cx asm5 -w30 –cs --eqx --secondary=no’ and visualized as dotplots with paf2dotplot (Jiang 2021). The resultant PAFs were analyzed with SyRI with the ‘--cigar’ option, and as synteny maps by plotting the SyRI calls with plotsr ((Goel and Schneeberger 2022).

Multi-genome synteny was calculated with ntSynt (Coombe *et al*. 2024) and visualized with ntSynt-viz ((Coombe, Warren and Birol 2025), using default parameters.

### Phylogenomics

#### PCA

75 Cannabis genome assemblies were downloaded from the Salk Institute Cannabis Pangenome website ((Lynch *et al*. 2025). Chromosome-scale scaffolds were purged of non-chromosomal contigs by filtering with ‘seqkit seq -m 10000000,’ to retain sequences over 10Mb. Unscaffolded assemblies were raised to chromosome scale, and to some degree purged of alternate contigs, by mounting to ‘Sour Diesel B’ chromosomes with ntJoin ((Coombe *et al*. 2020), using parameters ‘target_weight=1 reference_weights=’2’ no_cut=True overlap=False k=32 w=1000 G=100000’. Including the 2 Medcann and 8 Lighthouse haplotypes, a total of 85 10-chromosome drafts were then aligned against ‘Sour Diesel B’ with minimap2, using parameters ‘-ax asm5 -w30 --secondary=no --cs --eqx’, and the resultant BAMs called jointly with minipileup ((Li 2024) with parameters ‘-vc -a0 -s0 -q0 -Q0’.

The VCF output from minipileup was filtered for biallelic SNPs with vcftools (Danecek *et al*. 2011), using parameters ‘--min-alleles 2 --max-alleles 2 --max-missing 0.2 --maf 0.05 --remove-indels’, and pruned for linkage in PLINK2 using parameters ‘--indep-pairwise 50 10 0.1’. PCA was performed in PLINK2 with ‘--pca 84’. Cluster number (between 2 and 30) was evaluated via Elbow, Silhouette, Gap statistic, Calinski-Harabasz, Davies-Bouldin, and Dunn Index methods. Within the PCA, genotypes were clustered via k-means, Partitioning Around Medoids (PAM), or Gaussian Mixture Model (GMM) with k from 2 to 5, and visualized in R with ggplot2 (Wickham 2011).

#### Admixture

Linkage-pruned SNP calls were converted from VCF to BED with ‘plink2 --make-bed,’ analyzed in ADMIXTURE (Alexander and Lange 2011) with k from 2 to 5, and the results were then visualized in R with ggplot2. Results were truncated to three decimal places and then first sorted from bottom to top by percentage of ‘indica’ ancestry. Then, among 0% indica types, results were sorted from the top down in order of decreasing HEMP ancestry. Intermediate samples with neither ‘indica’ nor HEMP ancestry were sorted from bottom to top by decreasing ‘sativa’ fraction.

#### pan-NLRome

We ran NLR-Annotator (Steuernagel *et al*. 2020) on the complete assemblies to extract candidate loci with 1k flanking sequence, predicted proteins *ab initio* with AUGUSTUS (--species=arabidopsis --codingseq=on --genemodel=complete) (Hoff and Stanke 2019), filtered for length above 380aa ((Kourelis *et al*. 2021), and took as candidate proteins those sequences confirmed by hmmsearch (Eddy 1998) using our Cannabis NBS HMM (Pike, Kozik and Terán 2025) at stringency of 1e-10. The amino acid sequences were clustered with default parameters on the DIAMOND-DeepClust webserver ((Zimmermann *et al*. 2018; Buchfink *et al*. 2023), with a confirmatory analysis performed in OrthoFinder ((Emms and Kelly 2019). In a spreadsheet, we calculated presence-absence variation per orthogroup (Supp. Tables 7 and 8), and then elected to consider the CORE NLRome as those orthogroups occurring in 95% or more of genotypes, SHELL as those occurring in 15-95% of the population, and CLOUD as those found in less than 15%.

Data were processed on the ‘pyky’ node of the ZINE high performance computing cluster at the Pontificia Universidad Javeriana in Bogotá, Colombia. It contains 192 Xeon 8160 cores and 2Tb of RAM.

## Results

### Genomic DNA sequencing

Results are summarized in Supplemental Table 2. The 6 PromethION cells yielded from 35.6 to 79.8 Gb of sequence, with N_50_ ranging from 21.2 to 63.7 kb. Input coverage was 189x (ANC), 196x (HAW), 31x (SRI), or 91x (TRC). For Hawthorne OG, one PromethION cell was loaded with a library after treatment with a Short Read Eliminator XL kit (Circulomics, Baltimore, MD), and one with a library filtered above 50kb on the Blue Pippin. Yields were comparable (80 vs 66 Gb), while the fraction of reads above 50kb shifted from 8.4% to 63.1%.

Illumina coverage averaged 141 ± 18.4x for whole-genome shotgun (WGS) and 121 ± 13.5x for Hi-C (Supp. Table 2), prior to filtering.

### Assembly

Our process arrived at assemblies of 10 chromosome-scale pseudomolecules for the 8 haplotypes. The improvement derived from the several rounds of PECAT optimization is summarized in Supplemental Table 3. Relative to defaults, the modifications reduced the size of the primary haplotype draft from 1027 to 877Mb and the number of contigs from 546 to 262. They also increased the median contig (N_50_) from 3.8 to 12.0 Mb and the longest contig from 26 to 45 Mb. The fraction of BUSCO duplicates decreased from 23.7% to 12.4%, prior to purging.

Genome size, median contig length, and BUSCO scores are shown in Table 1.

**Table 1.** Essential statistics for the eight assemblies.

| genotype | Anders CBD |  | Hawthorne OG |  | Sri Lankan |  | Travis CBD |  |
| --- | --- | --- | --- | --- | --- | --- | --- | --- |
| haplotype | a | b | a | b | a | b | a | b |
| scaffolds | 10 | 10 | 10 | 10 | 10 | 10 | 10 | 10 |
| contigs | 122 | 133 | 129 | 173 | 444 | 525 | 207 | 181 |
| size (Mb) | 748 | 713 | 708 | 678 | 646 | 642 | 735 | 739 |
| $N_{50}$ (Mb) | 11.0 | 11.3 | 11.4 | 6.3 | 2.8 | 2.0 | 7.9 | 8.2 |
| BUSCO | 96.48% | 95.35% | 96.65% | 95.91% | 89.72% | 88.35% | 97.55% | 97.63% |
| single | 90.89% | 90.15% | 93.81% | 93.85% | 87.27% | 85.60% | 94.50% | 94.02% |
| duplicated | 5.59% | 5.20% | 2.84% | 2.06% | 2.45% | 2.75% | 3.05% | 3.61% |
| fragmented | 0.17% | 0.04% | 0.34% | 0.52% | 0.13% | 0.26% | 0.00% | 0.13% |
| incomplete | 0.00% | 0.00% | 0.00% | 0.00% | 0.00% | 0.00% | 0.00% | 0.00% |
| missing | 3.35% | 4.60% | 3.01% | 3.57% | 10.15% | 11.39% | 2.45% | 2.24% |

Changes in Quality Values (QV) with each round of polishing, as calculated by ‘yak qv’ via comparison to short read 21mers, are shown in Supplemental Table 4. On average, raw reads of QV 16.04 were assembled and polished to QV 24.71.

### Alignment

Dotplots of the 8 haplotypes, aligned to ‘Sour Diesel B,’ are shown in Figure 1. Overall collinearity was quite good, with the exception of a large inversion on chromosome 3, which we analyze below. Some gaps may be seen, which we ascribe to incomplete phasing.

**Figure 1.**
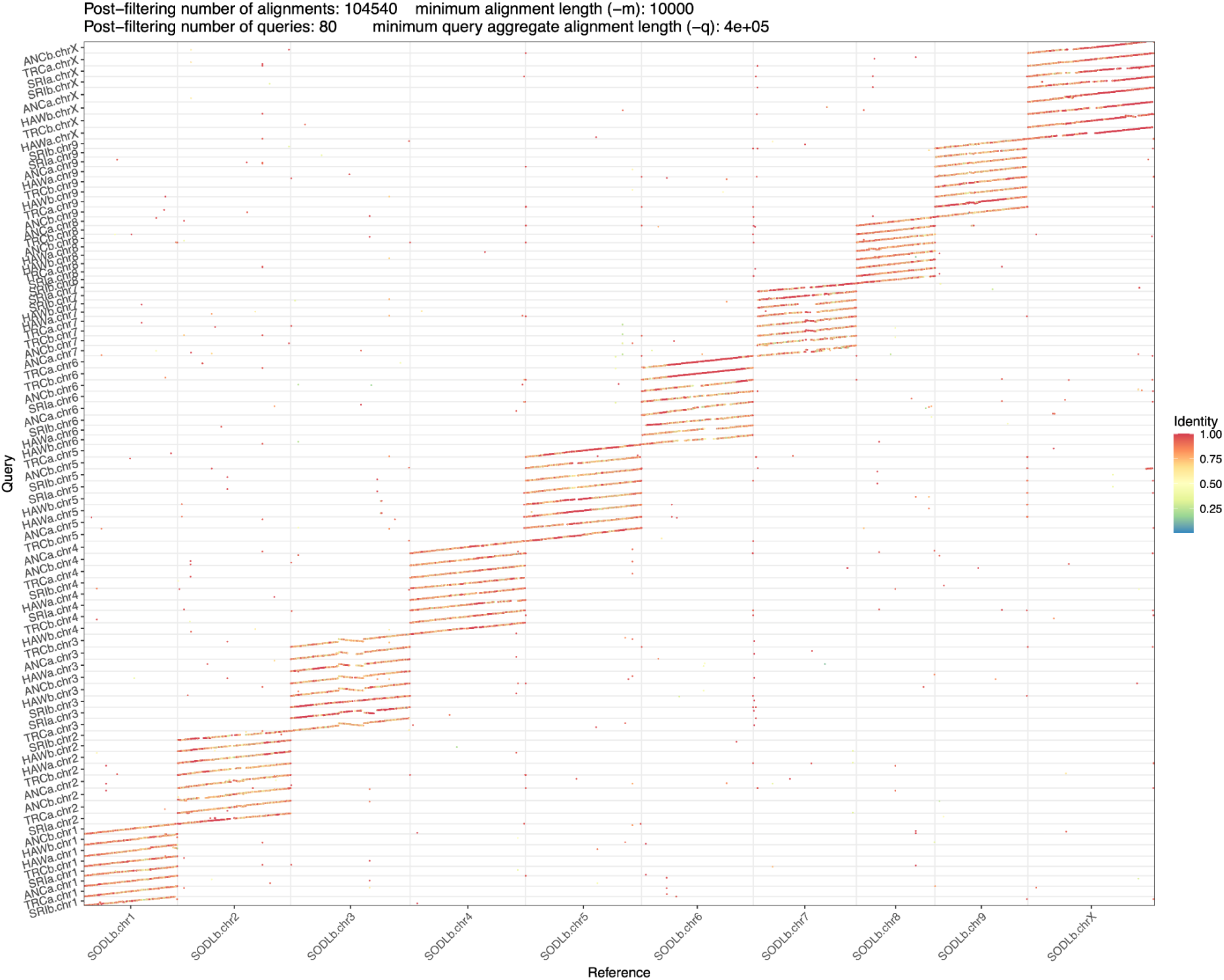
8 Cannabis haplotypes arranged against ‘Sour Diesel B’.

We visualize macrosynteny, calculated from kmers common to all drafts, with ntSynt-viz (Figure 2). We include PR and CP, ‘Sour Diesel B’ (SODLb), and the previous and current NCBI references, cs10 ((Grassa *et al*. 2021) and Pink Pepper ((Lim 2023).

**Figure 2.**
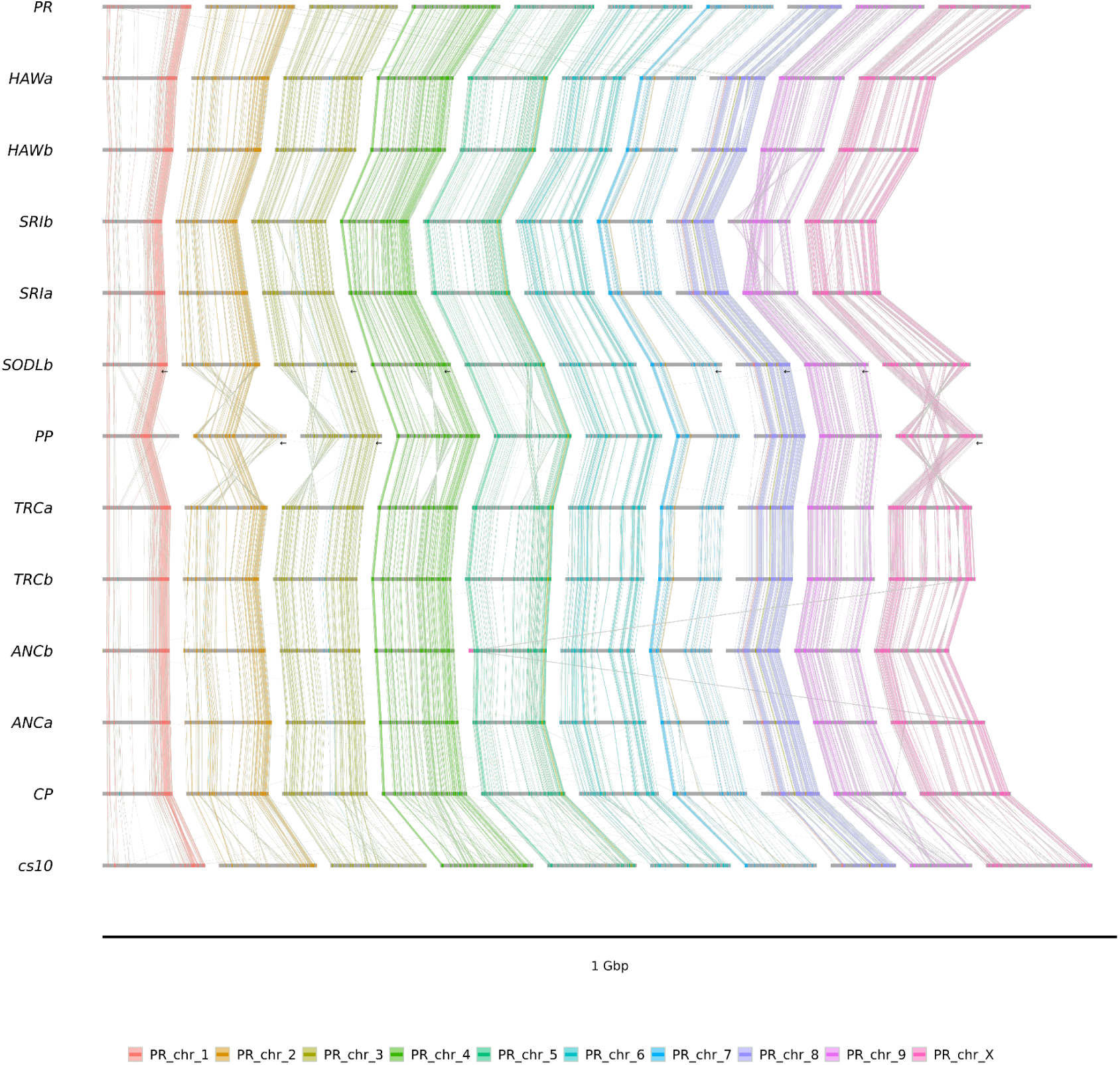
Visualizations of minimizers common to the 8 Lighthouse haplotypes, PR and CP, Sour Diesel B (SODLb), and the current and former NCBI references Pink Pepper (PP) and cs10. Arrows indicate reverse-complementation.

Relative to SODLb, SyRI reported an average of synteny of 578Mb, 37 Mb of inversions, 16 Mb of translocations, 5.2 Mb of duplications, and 65.2 Mb unalignable. Within syntenic regions, on average, 2.64M SNPs, 289k insertions, and 365k deletions were called, and an average of 328Mb per haplotype was called as Highly Diverged Regions (HDR). Across the 8 assemblies, inclusive of HDR, we identified 141,382 variants over 10kb, 10,226 over 100kb, and 134 over 1Mb. Complete SyRI results for structural and sequence variation are included in Supplemental Tables 5 and 6.

To clarify the nature of a large inversion seen in chr3 when aligning to SODLb, we visualized alignment of a representative set of unphased dual contigs to it (Supp. Fig. 1a), and also decomposed the SODLb chromosome into its component contigs (Supp. Fig 1b).

### Annotation

Annotation counts for all genes, NLRs, TPS, and repeat content and LAI, are summarized in Table 2:

**Table 2.** Annotation counts for cs10, which we used as a source for Liftoff, and the 8 Lighthouse haplotypes. Repeat content for cs10 is as reported previously (Grassa *et al*. 2021).

|  | cs10 | ANCa | ANCb | HAWa | HAWb | SRIa | SRIb | TRCa | TRCb |
| --- | --- | --- | --- | --- | --- | --- | --- | --- | --- |
| gene | 29489 | 29359 | 29048 | 28821 | 28552 | 26925 | 26753 | 29098 | 29335 |
| gene (%) | 100.0% | 99.6% | 98.5% | 97.7% | 96.8% | 91.3% | 90.7% | 98.7% | 99.5% |
| mRNA | 33382 | 32396 | 32018 | 31965 | 31532 | 30006 | 29660 | 32087 | 34423 |
| CDS | 33382 | 32350 | 31938 | 31904 | 31455 | 29908 | 29557 | 32058 | 32381 |
| lncRNA | 3041 | 3544 | 3590 | 3429 | 3378 | 3085 | 3106 | 3564 | 3616 |
| rRNA | 12 | 14 | 22 | 18 | 10 | 11 | 41 | 31 | 40 |
| snoRNA | 1576 | 1590 | 1554 | 1533 | 1557 | 1405 | 1433 | 1546 | 1536 |
| snRNA | 99 | 88 | 87 | 87 | 92 | 90 | 82 | 93 | 94 |
| NLR-Annotator | 259 | 264 | 253 | 253 | 255 | 244 | 246 | 269 | 287 |
| NBS HMM | 295 | 301 | 296 | 292 | 294 | 270 | 283 | 315 | 339 |
| NLR merged | 244 | 251 | 238 | 239 | 242 | 232 | 234 | 255 | 274 |
| TPS | 47 | 48 | 49 | 39 | 41 | 41 | 46 | 43 | 47 |
| repeats | 63% | 74.89% | 74.39% | 74.18% | 73.96% | 74.52% | 74.62% | 75.29% | 75.32% |
| LAI | - | 12.66 | 13.64 | 18.99 | 18.79 | 17.81 | 17.41 | 14.47 | 14.94 |

### Phylogenomics

The population structure, as revealed by linkage-pruned biallelic SNPs, was analyzed with PLINK2 and visualized by PCA in R with ggplot2 (Fig. 3). The six methods used to determine subpopulation number (k) produced inconsistent and highly variable results, ranging from 11 (Elbow method) to 30 (Davies-Bouldin and Gap statistic) (Supp. Fig. 2). Therefore, to arrive at a useful classification scheme, we evaluated k from 2 to 5, based on the well-defined divisions and known pedigrees of HEMP, MJ, and HC_HEMP, and as corroborated by previously published reports (Lynch *et al*. 2016; Grassa *et al*. 2021; Jin *et al*. 2021; Chen *et al*. 2022; Wang *et al*. 2025). We found that the Partitioning Around Medoids (PAM) (Supp. Fig. 3) and Gaussian (Supp. Fig. 4) methods universally failed to maintain MJ or HEMP as monophyletic groups, as did a variety of dendrogram-based methods (data not shown). Only k-means, with k=4, faithfully grouped genotypes in accordance with their established use cases, which are consistently supported by the known pedigree data (Fig. 3) (Lynch *et al*. 2025). The one exception is Durban Poison (DP), an MJ type from Durban, South Africa, which is typically considered to be a pure ‘sativa’ landrace, and which here is called as HC_HEMP. K-means clustering with k of 2, 3, or 5 suffered from errors similar to the other methods. (Supp. Fig. 5.)

**Figure 3.**
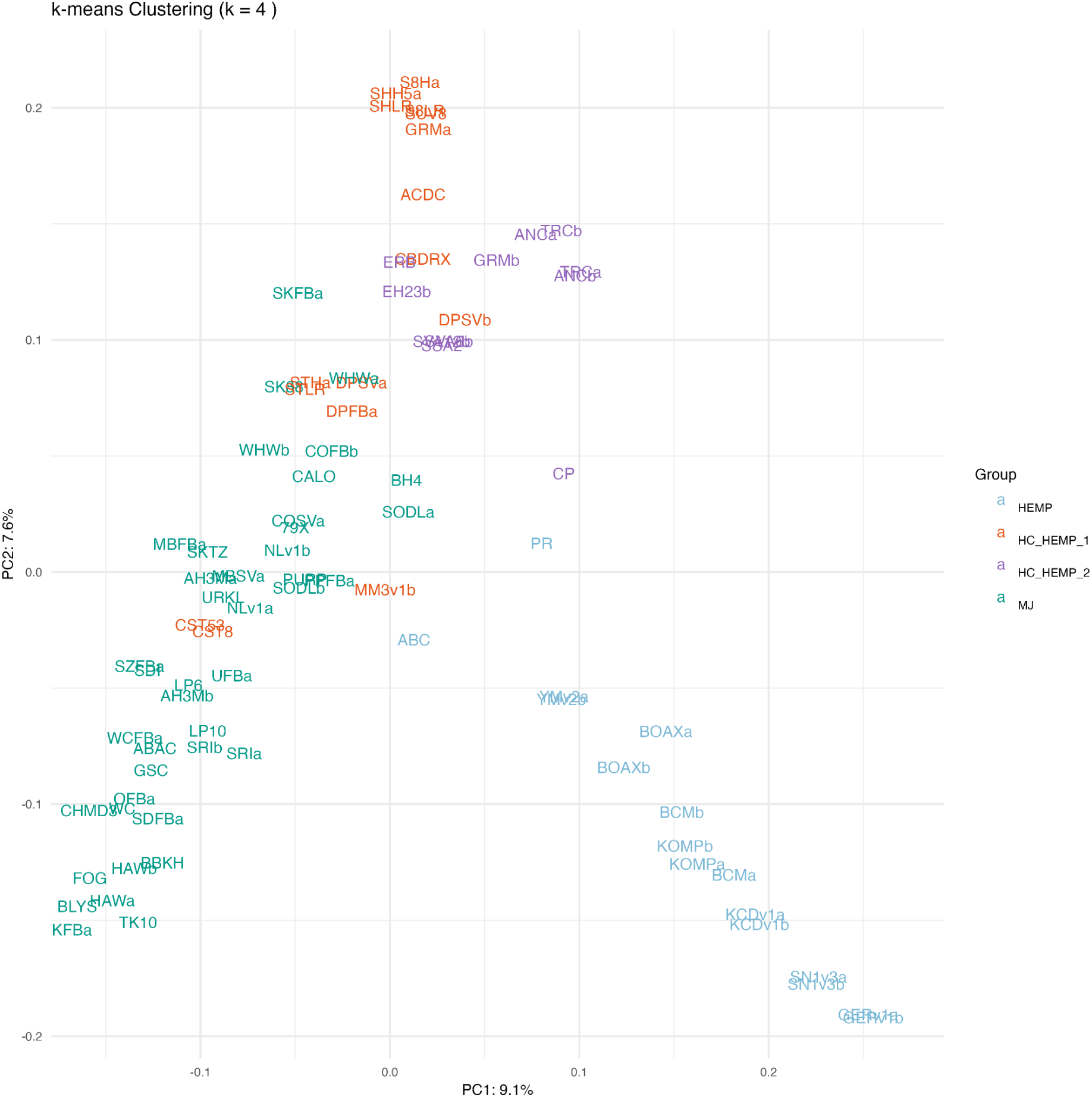
PCA of 85 Cannabis genome assemblies with k=4. Genotype codes are listed and described in Supplemental Table 1. Blue codes indicate HEMP, purple and red both HC_HEMP, and green MJ genotypes.

**Figure 4.**
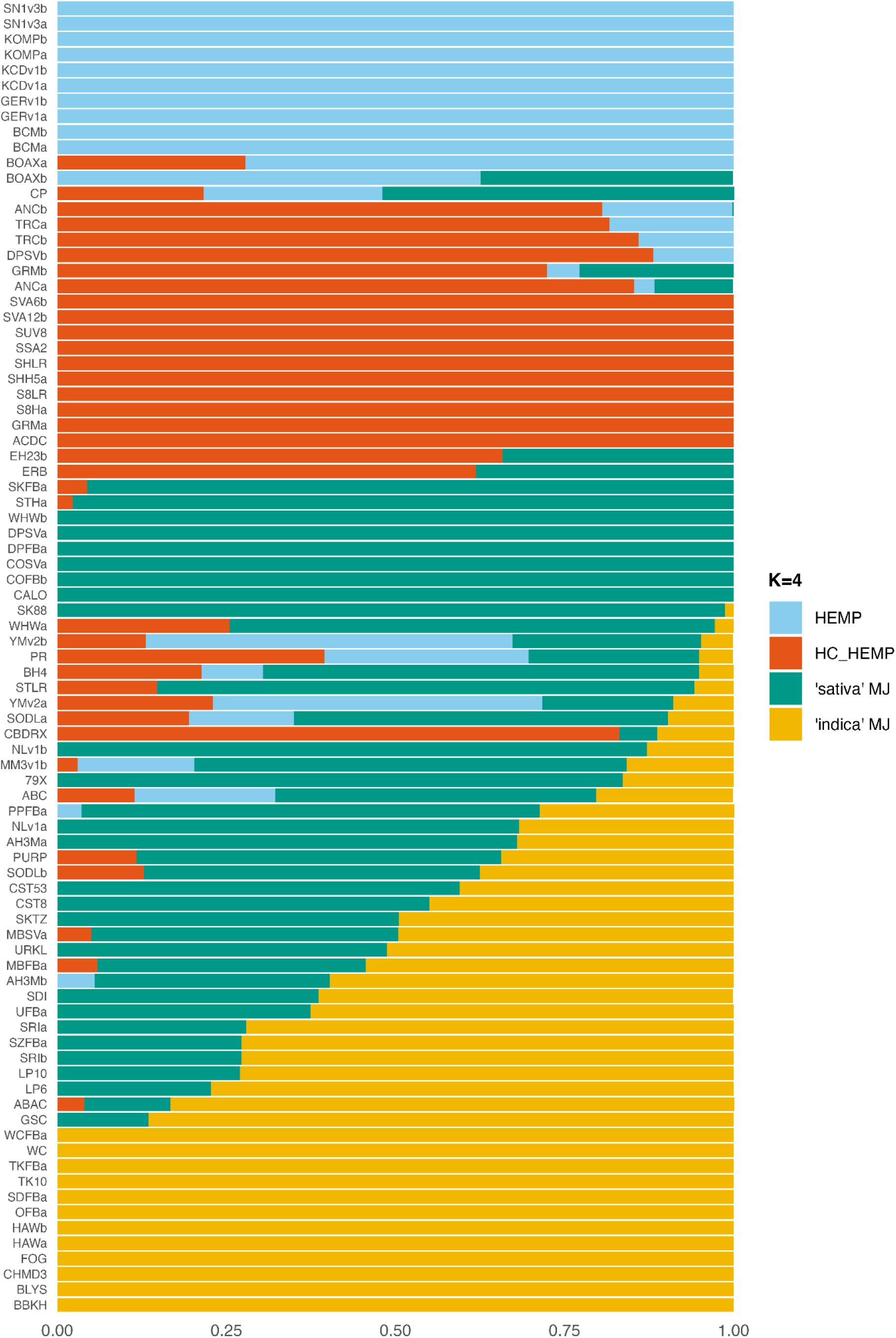
Admixture analysis of 85 Cannabis genomes with k=4. Genotype codes are listed and described in Supplemental Table 1.

Ancestry estimates were calculated with ADMIXTURE from the same data and visualized as a bar plot in Figure 4. Again, we found that k=4 maintained the known distinction between MJ and HC_HEMP in a way that k of 2, 3, or 5 did not (Supp. Fig. 6).

In both cases, k=4 provides clean distinctions between major groups that are coherent with our prior knowledge of the relevant pedigrees and use cases. K-means analysis of the PCA divided HC_HEMP into 2 groups, one with HEMP-admixed individuals (HC_HEMP_2) and one without (HC_HEMP_1). However, ADMIXTURE finds a single HC_HEMP group and divides MJ into two groups, which correlate well with grey-market perceptions of ‘sativa’ and ‘indica’ heritage, with ‘indica’ representing dense, resinous, fast flowering ‘hashplant’ types from Southwest Asia, and ‘sativa’ representing taller, less potent, longer flowering lineages from Latin America, Africa, and South and Southeast Asia (McPartland 2017).

#### pan-NLRome

NLR proteins were predicted and clustered into orthogroups. From 25,963 NLR-Annotator loci, 10,216 proteins were predicted by AUGUSTUS, of which 7,933 were further filtered using the Cannabis NBS HMM profile. The putative resistance proteins were aligned with DIAMOND and then clustered into orthogroups by DeepClust or OrthoFinder. Distribution among CORE, SHELL, and CLOUD, which are present in >95%, 15-95%, or <15% of genotypes respectively, are shown in Table 3, with genotypes per orthogroup listed in Supplemental Table 7 (DeepClust) and 8 (OrthoFinder).

**Table 3.** Distribution of predicted NLR proteins among CORE, SHELL, and CLOUD classes according to the prevalence (‘prev.’) of the orthogroup (OG) across the 85 cannabis genomes..

| class | prev. | DeepClust |  |  | OrthoFinder |  |  |
| --- | --- | --- | --- | --- | --- | --- | --- |
|  |  | OG | NLR | NLR % | OG | NLR | NLR % |
| CORE | >95% | 34 | 5225 | 65.86% | 57 | 4794 | 64.84% |
| SHELL | 15-95% | 20 | 1605 | 20.23% | 40 | 1997 | 27.01% |
| CLOUD | <15% | 42 | 1103 | 13.90% | 110 | 603 | 8.16% |
| TOTAL |  | 96 | 7933 |  | 207 | 7394 |  |

## Discussion

### Technical considerations: optimizing PECAT output

The Phased Error Correction and Assembly Tool (PECAT) pipes the output from a minimap2 all-versus-all alignment to custom scripts, first for error correction and then to generate a pseudohaploid draft. Corrected reads are then mapped to the pseudohaploid, and SNPs are called with Clair3 and phased with WhatsHap. Inconsistent overlaps, i.e., those that induce phase switches, are eliminated from the initial string graph, which is then newly assembled to give a set of dual contigs. We found that the stock configuration for Arabidopsis did not adequately collapse the haplotypes, leading to oversized drafts with BUSCO duplication over 20% (Supp. Table 3).

By increasing Minimap2’s index and PECAT’s block size from 4Gb to 400 Gb, all the input reads were processed in one go, in true ‘all-versus-all’ fashion. Besides, by avoiding the need to concatenate subsets prior to overlap filtering, temporary disk requirement was reduced from several Tb to less than 1 Tb. With about 90 Gbp of input sequence, RAM use peaked between 650 and 700 Gb (data not shown).

PECAT’s default configuration extracted the longest 80x of reads for correction, typically leading to assembly input of about 65x (data not shown). We found that increasing the input to correct the longest 112x, and assemble the longest 88x of corrected reads, created fewer and longer contigs (Supp. Table 3). We note that SRI and TRC began with less coverage than this (31x and 91x), so that assembly for them began with lower coverage (24x and 86x). As per the default PECAT configuration for human and bovine genomes, we used -f 0.005 for correction and -f 0.001 for assembly, which discards the 0.5% and 0.1% most abundant kmers. This prevents the formation of massive temporary files, and did not much change the assembly statistics relative to the Minimap2 default of 0.02% (Supp. Table 3). As well, we found that correcting and assembling with 19mers yielded greater contiguity than correcting with 17mers and assembling with 19mers (data not shown). We were unable to test the default, which corrects with 15mers, due to its creating more than 3.2 Tb of temporary overlap files. We suspect that correcting from these much more stringent overlaps contributed to the creation of longer contigs with lower QV.

The stock configuration filters long overhangs, to reduce the risk of chimerism. We retained the filter to require no more than 0.3 of the read in overhang, and eliminated the hard limit of 3000 bp. Because the data used by Nie et al. (2024) had N50 of 12k, these parameters were similar, but with our ultra-long libraries, which still require at least 0.5 of the read to be in overlap, we found that allowing longer overhangs permits additional contiguity to be discovered (Supp. Table 3).

To increase haplotype collapse, we raised -r from 500 bp to 20 kbp, which broadens the gap across which minimap2 can chain minimizers, and therefore extends alignments across an indel or other SV. This brought the size of the pseudohaploid draft much closer to the genome size predicted by flow cytometry (808 Mb for female genotypes) (Gao *et al*. 2020), and reduced BUSCO duplicates after diploidization.

To phase SNPs, PECAT deploys Clair3 as well as a native method called FSA. We found that FSA increased BUSCO duplicates and so relied exclusively on Clair3. For phasing, we used corrected reads, with the expectation that SNP calls would be more precise, and we adjusted the minimap2 parameters to use the Range-Min-Query (RMQ) mode, as it is better at aligning across long indels. We used an initial bandwidth of 1kb and 250kb for extension, after evaluating the size distribution of SVs found between the trio-binned haplotypes of Punto Rojo and Cherry Pie (Pike, Kozik and Terán 2025), where we found 440 SVs over 100kb and only 39 over 250kb. We also preclude secondary mappings, as is common in phasing pipelines (Porubsky *et al*. 2021; English *et al*. 2022).

The Sri Lankan genotype, which was collected at a preliminary stage of the investigation, did not benefit from electrophoretic size selection, and was also sequenced to much lower depth. Therefore, its haplotypes are much less contiguous and complete than the others. The N50 of 2.8 and 2.0 Mb certainly stems from a lack of sufficient ultra-long coverage. The BUSCO scores of 89.7% and 88.4% are a bit disappointing, but we note that an analysis of the two haplotypes together gives a BUSCO score of 98.7%, indicating that the dual assembly has high completeness, but is incompletely phased. We were not able to remedy this situation with tools such as Quickmerge, ntJoin, or RagTag. We note that Sri Lankan’s genic and NLR counts also lag by about 10%, and so we acknowledge that there will be some false negatives in its downstream analysis.

Finally, we acknowledge the recent release of HERRO (Kang *et al*. 2023), which is theoretically able to correct ONT R9 reads to Q20 or higher, through the use of a neural net trained on the pore’s specific error profile. In a test, we found that HERRO-corrected reads assembled into a draft that had 45% lower N_50_ (6.5 Mb vs 11.8 Mb) and only fractionally higher QV (18.9 vs 18.1), and so we relied exclusively on the older method of haplotype-aware error correction contained within PECAT.

### Diploid polishing

A challenge in diploid assembly is to reduce sequencing errors without re-introducing previously phased SNPs. These assemblies have QV lower than what is typical for modern plant genomes, presumably in part due to the subtractive nature of PECAT’s diploidization process, and in part because they have been polished carefully with the aim of not inducing per-base phase switches, known as Hamming errors (Hamming 1950).

PECAT polishes the drafts by re-using the phasing information to only map reads specific to each haplotype. When compared to short read kmers, this raises base level precision from about 98.2 to 98.5%. To reduce cross-haplotype alignments, we mapped Illumina reads to the concatenated primary + alternate (pri-alt) assembly after purging haplotigs from each. We called 1/1 variants with Clair3 and repeated the procedure, which raised precision above 99% or Q20 (Table 3). Finally, the contigs were polished four times with ntEdit, which uses a kmer database to locate errors and replace them with the most plausible alternative (Warren *et al*. 2019). Because this approach is not read based, it should not add switch errors, as both alleles will be present in the short read kmer table and, therefore, neither will be called as a potential error. However, because ntEdit considers both the ‘leading’ and ‘trailing’ kmer when making a substitution, and only substitutes when both are present in the short-read table, it is unable to correct errors when two are found separated by fewer than 2k bases.

The quality (QV) of these assemblies averages 24.71 (Supp. Table 4), implying base-level accuracy of 99.66%. While this pales in comparison to recent HiFi assemblies scored at QVs of 50 or higher, as the contiguity and completeness are comparable, and the overall synteny is high, this does not preclude a comparative analysis.

### Scaffolding

The Hi-C method of scaffolding arranges contigs in three main phases: grouping, ordering and orientation. The high repeat content of plant genomes presents challenges for each. Preliminary trials with several algorithms produced ‘megascaffolds’, with some or all chromosomes conjoined, a known phenomenon (Sur and Myler 2022). As well, some chromosomes remained fragmented in 10 or more scaffolds, presumably due to ambiguity in repetitive regions. We found that scaffolding on a per-chromosome basis resolves both issues.

We made use of the pseudohaploid intermediate produced by PECAT to create a chromosome-scale draft to guide the diploid assemblies. To avoid over-grouping, we combined the primary and alternate pseudohaploid contigs and binned them into linkage groups, after aligning them to ‘Sour Diesel B’ (SODLb), the most contiguous haplotype from the Salk Institue Cannabis Pangenome (Lynch *et al*. 2025). Each contig was primarily aligned to one SODLb chromosome. This intermediate structure, scaffolded to a pseudomolecule by HapHiC, is oversized, due to the inclusion of alternate haplotigs, but no large interchromosomal translocations were observed.

To sort the diploid contigs, we first mapped them to the pseudohaploid pseudomolecules, to bin them per chromosome. The native pseudohaploid was preferred, as we observed that some diploid contigs would bin differently with the use of different heterologous references. Next, each linkage group was scaffolded with GreenHill (Ouchi, Kajitani and Itoh 2023), which is a recently published algorithm derived from the strategy of Falcon-PHASE (Kronenberg *et al*. 2021), and not previously demonstrated capable to scaffold plant genomes. GreenHill identifies allelic contigs and collapses them to produce consensus contigs, prior to ordering and orienting based on Hi-C as well as long-read mappings. Finally, the collapsed contigs are diploidized based on stored SNP information. Aligning these scaffolds to SODLb reveals 10 pairs of chromosome-scale pseudomolecules.

However, the Greenhill scaffolds were inadequate for three reasons: many orientation errors were evident, length polymorphisms were necessarily pruned to create bubble edges of equal length, and many SVs likely failed to be reconstituted when said bubbles were ‘unzipped’. Therefore, this data structure may be best considered as a linear database of haplotype-specific kmers, which we next used to bin the diploid contigs via alignment. The hap0 and hap1 scaffold sets from GreenHill were concatenated and the diploid contigs mapped to them. Next, the putative haploid contig sets were each mapped to SODLb, and some duplicate haplotigs that survived the first purge were either removed or relocated to the other haplotype, based on a visual inspection. We presume that this second purging could and should be automated if the pipeline were to be used more widely. Finally, each set of haplotigs was scaffolded separately with YaHS, which deploys a sophisticated algorithm allowing to resolve ambiguous orientations and avoid the formation of biallelic ‘hairpins’.

### Alignment

When the Lighthouse haplotypes are aligned against SODLb (Fig. 1), no obvious misassemblies are observed, which accounts for the high quality of our assembly and scaffolding workflow. We found a large (∼22Mb) inversion in the center of chr3, which we presume includes the centromere, and which aligns perfectly with contig boundaries (Supp. Fig 1b). Therefore, we infer that a 4-contig scaffold was placed in the correct order but with incorrect orientation, a previously reported issue (Ghurye *et al*. 2019) with the 3D-DNA algorithm, which was used by Lynch et al. (2025) to arrange contigs into scaffolds from Hi-C data. In our trials, we found that YaHS provided more accurate results, as corroborated by two recent reports that evaluated different scaffolding tools in Arabidopsis, rice and strawberry (Hou, Wang and Pan 2023; Obinu, Trivedi and Porceddu 2024).

Aligning a dual set of contigs assembled from ONT ultra-long reads against the HiFi-based SODLb chr3 is instructive, as contig extension with the two platforms appears to fail differentially. One may note that ONT contigs 4, 18, 33, and 3_1 extend well into the inversion, from both sides in both haplotypes, but do not support it (Supp. Fig. 1a). However, the homologous HiFi contig (ctg5, Supp Fig. 1b) spans the gap between them.

On chr7, spanning the B locus, we observe a lack of linearity (Fig. 1), or a blank space with no common minimizers (Fig. 2). This is not unexpected, as these genomes include both B_D_ and B_T_ karyotypes, i.e. the loci for the CBD and THC synthases (de Meijer *et al*. 2003), which each lie in massive haploblocks that appear to not recombine (Grassa *et al*. 2021).

When synteny is visualized, in terms of common minimizers across the assemblies, a common architecture is apparent across all genomes (Fig. 2). However, it is worth noting that this architecture is not shared by the current NCBI reference, Pink Pepper, which was created without benefit of the most modern assembly and scaffolding algorithms. Even at this scale, it is clear to see the difference among ONT assemblies created with scant or abundant ultra-long coverage. The former includes PR, CP, SRIa, SRIb, and cs10, all of which display unique small rearrangements that surely are technical errors. However, SRI shows fewer interchromosomal translocations than PR and CP, despite emerging from similar readsets that produce similar contig N_50_. This highlights the added value of scaffolding with Hi-C, rather than by arranging contigs to a reference.

Among the other 6 Lighthouse drafts (ANC, HAW, and TRC), observed macrosynteny is well conserved, with just one glaring disagreement: a piece of ANCb chr5 which lies in chrX in all other drafts. While this may be a legitimate translocation, and is well supported by both long-read and Hi-C coverage, we suspect it is more likely to be an assembly error.

### Genome annotation

Annotations are summarized in Table 2. As shown previously (Pike, Kozik and Terán 2025), Liftoff (Shumate and Salzberg 2021) seems competent to recover nearly all reference genes in draft genomes, without needing to mask repeat elements *a priori*. Here, the six haplotypes built from high-coverage ultra-long libraries have 95.8 −99.5% gene content of the cs10 reference. The two Sri Lankan drafts score slightly lower, with 90.7% and 91.3%, consistent with their BUSCO scores, that also suffer from inadequate diploidization, mainly due to shorter reads and lower sequencing depth.

With respect to NLR resistance genes, the counts reported here range from 229-272 and are comparable with the 244 found in cs10 and the 227 and 240 we reported previously for PR and CP, respectively (Pike, Kozik and Terán 2025).

The TPS counts range from 40 to 49, compared with 47 in cs10. We acknowledge that sequence homology-based prediction of TPS products is inaccurate, and that functional characterization via cloning, or genome editing, is necessary to validate and quantify their activity (Booth *et al*. 2020). Of particular relevance for cannabis breeding, putative genes coding for TPS participating in myrcene or limonene synthesis are frequently responsible for producing these, as well as other monoterpenes, in a fixed ratio (Booth, Page and Bohlmann 2017). Therefore, we suggest these results could be best taken as a preliminary genome-wide scan to identify SVs of interest, related to cannabis ‘terptypes,’ for further inquiry.

Regarding genome wide repeats, repeat content ranged from 73.96-75.32%, and the LTR assembly index (LAI) ranged from 12.66-18.99 (Table 2). Here, the use of panEDTA, which develops a common repeat library across all samples, raised the repeat content by 2-4% relative to EDTA (Ou *et al*. 2019), primarily due to an increase in Copia/Ty1 content (data not shown).

### Phylogenomics and Population Structure

#### PCA

PCA revealed a population structure similar to that reported previously for region-of-origin cannabis landraces (Ren *et al*. 2021), with PC1 showing strong differentiation between ‘drug-type’ and ‘hemp-type’ (Fig. 2), which here we call marijuana (MJ) and hemp (HEMP), as do Lynch et al (2025). Further, most high-cannabinoid hemp (HC_HEMP) types were located in-between the former differentiated clusters, similarly as was found and reported as ‘basal cannabis’ and ‘drug-type feral’ in the analysis of Ren et al. This illustrates a pattern where modern hybridization of MJ with HEMP seemingly could have induced a recovery of genotypes similar to the natural form of the plant, i.e., prior to its improvement by humans.

PC1, which explained 9.1% of variance, appears to correlate with cannabinoid content, which is known to be heritable (Hennink 1994), polygenic (Woods *et al*. 2021), and the target of diversifying selection among marijuana and hemp breeders. PC2, which explained 7.6% of variance, is less easily interpretable, as it shows substantial and continuous variance within both MJ and HEMP, with the most potent hashplants and most fibrous hemp types occupying similar positions along it (Fig. 3).

HC_HEMP genotypes appear to primarily cluster at the intersection of two axes that run skew to PC1 and PC2. One, the ‘marijuana axis’, extended from ‘indica’ types such as Bubba Kush (BBKH), Hawthorne OG (HAW) and Triangle Kush (TK6, TK10), through a hybrid zone including the widespread Sour Diesel (SODL) and Girl Scout Cookies (GSC) cultivars, to ‘sativa’ types such as Durban Poison (DPFB, DPSV), Skunk (SK88, SKFB), and White Widow (WHW). Here, we use ‘indica’ and ‘sativa’ in their vernacular sense, following McPartland ((McPartland 2017). We acknowledge that this dataset lacks pure ‘Haze’ varieties, such as Neville’s Haze or Oldtimer’s Haze, which are rare in the market due to their long life-cycle times and relatively low cannabinoid content, and which we speculate might lie in-between the Skunk-Durban-White Widow boundary and the HC_HEMP group.

The orthogonal hemp axis groups, at its far end, types used exclusively for fiber, such as Kompolti (KOMP) and Santhica (SN3v1), while HC_HEMP types BoAx (BOAX) and Abacus (ABC) lie closer to the marijuana group.

With respect to clustering methods, while k-means provides useful and meaningful clusters in practice, and outperforms other methods (Supp. Figs. 3 and 4), we should note some simplifying assumptions within the algorithm that may cause errors with a real-world dataset. K-means assumes that each group has equal membership, and equal variance, and is spherical in shape. Here, our a priori knowledge of the input genotypes suggest that we have 12 HEMP, 26 HC_HEMP, and 47 MJ. As well, the PCA suggests that HC_HEMP contains less variance than HEMP or MJ, and that the two latter groups may be more oblate than spherical. Future work along these lines might seek to eliminate redundant MJ genotypes, and more widely sample HEMP, to reduce the possibility of spurious aggrupations. Fortunately, it has been shown that k-means is not very sensitive to deviations from these presumptions (Craen *et al*. 2006), which is consistent with the good results presented here.

We found that typical statistical methods recommended unusably high values of k (Supp. Fig. 2), and that k=4 accurately classed known types better than k of 2, 3 or 5 (Supp. Fig 5). Group 1 (red) clusters exclusively HC_HEMP, and includes our drafts of Cherry Pie (CP), Anders CBD (ANC), and Travis CBD (TRC). Group 2 (green) englobes exclusively MJ, and includes our Sri Lankan (SRI) and Hawthorne OG (HAW) assemblies. HAW lies close to putative relatives such as Fatso OG (FOG) and BBKH, while SRI, an exotic landrace, lands in-between GSC and Lemon Party #10 (LP10). Group 3 (purple) clusters also exclusively HC_HEMP; however, several of its members appear dispersed among MJ group: Sour Tsunami (STLR), Sour Tsunami x HO40 (STHa), Chem4 x Sour Tsunami F1 (CST53), and Chem4 x Sour Tsunami F2 (CST8). Sour Tsunami enjoys a reputation as one of the most flavorful and potent HC_HEMP types, and the gray literature offers a simple explanation for its placement within our PCA: it is reported as a hybrid between AC/DC (ACDC) and Sour Diesel (SODL), repeatedly backcrossed to SODL, with selection for the high CBD, low THC chemotype. Here, ACDC also locates within Group 3, but lies in the distinct HC_HEMP cluster at top, while the Sour Tsunami varieties cluster closer to MJ types SODLa and SODLb. This corroborates the underground pedigree, interestingly and tightly linked with main cannabis uses and breeding strategies, which definitely highlights the discriminatory power of our PCA.

On the other hand, group 4 (blue) clusters mostly HEMP, with the inclusion of ABC and BOAX, which are HC_HEMP, and the Colombian landrace Punto Rojo (PR), which is MJ. Remarkably, PR is the only high THC, low CBD chemotype that is not classified as MJ in this scheme.

#### ADMIXTURE

ADMIXTURE analysis of the same callset reveals somewhat different groupings. Here, the blue HEMP and purple HC_HEMP groups appear similar to those found by k-means clustering, with just one HC_HEMP group and not two, but now many MJ samples appear to be hybrids. We interpret this as representing various fractions of ‘sativa’ and ‘indica’ ancestry, with ‘indica’ types deriving primarily from Afghani landraces ((McPartland 2017), and ‘sativa’ describing landrace parentals from a vast swath of terrain that extends from SE Asia across India, to southern Africa and Latin America.

Reducing the complexity in this manner also reveals insights relevant to breeding, for example among HC_HEMP types that do not fall cleanly into that group. Our phased Anders CBD assembly has primarily HC_HEMP ancestry, but ANCa appears to include some HEMP and some ‘sativa’ ancestries, while ANCb includes only a larger fraction of HEMP. Both haplotypes of Travis CBD appear similar to ANCb. Cherry Pie, meanwhile, shows more HEMP content than any other HC_HEMP, with the exception of BoAx. It was also unexpected in showing about 50% ‘sativa’ ancestry. We suggest that the presence of some HEMP ancestry could then be relevant to compliance with strict THC limits in legislation, whether it be harvesting before THC rises over 0.3%, as in the US, or the Colombian standard of <1.00% THC at maturity, as determined by government agronomists. Cherry Pie is unusual in its HEMP content, and, in field trials, was the only one of 8 North American HC_HEMP accessions that could satisfy the stringent Colombian metric.

Near the middle of the spectrum, a cluster of 7 genotypes show unusual ancestry admixtures with both HEMP and ‘indica’ ancestry: Blueberry Haze (BH4), SODLa, STLR, CBDRx, both haplotypes of Yunma (YMv2), and Punto Rojo. Blueberry Haze, an MJ type, is the only named ‘Haze’ in this dataset, reported as a hybrid of it with Blueberry, a well-known ‘sativa’-’indica’ hybrid created in the state of Oregon in the 1970s (Short 2019). Therefore, this placement could be taken as scant corroboration that ‘Haze’ varieties may include some HEMP, and we restate that this dataset would be more comprehensive with the inclusion of additional ‘Haze’ genotypes.

We note in passing that SODLa, but not SODLb, includes some HEMP ancestry, which renders its admixture rather similar to that of Punto Rojo, albeit with more ‘sativa’ and ‘indica’ and less HEMP and HC_HEMP. This is surprising as SODL is perhaps the most relevant genetic in modern MJ, and is known for its high potency and suitability for indoor cultivation. While again admittedly scant, we imagine that investigating the nature of the apparent HEMP introgression in top-shelf MJ types, such as BH4 and SODL, may reveal some unusual allelic novelties.

With regards to Punto Rojo, it is remarkable that the genetic admixture to which it is most similar is Yunma, an open-pollinated dual-purpose (fiber and seed) variety from China, specifically the SW province of Yunnan, adjacent to Burma, which falls within the range where paleopalynology suggests Cannabis diverged from Humulus about 27 Mya (Zhang *et al*. 2018; McPartland, Hegman and Long 2019). The two haplotypes of YMv2 appear very similar, and PR lands between them, due to our sorting by decreasing fraction of ‘indica’ ancestry. We hypothesize that this pattern does not reflect strong selection for cannabinoid content, as is common among less admixed types, and instead derives from centuries of adaptation to cultivation, in tropical to subtropical montane environments, prior to the invention of agrichemicals.

The pedigree of PR is murky, and so we cannot rule out *a priori* the possibility that it may be a modern hybrid of HEMP, HC_HEMP, ‘indica’, and ‘sativa’. However, its morphology and chemotype are highly consistent with anecdotal reports of classical Colombian landraces, and so we proceed on the assumption it is a representative example of them.

Finally, across PCA and ADMIXTURE, both estimates derived from whole genome alignments, we find that Punto Rojo is positioned near the center of the genomic spectra found within Cannabis. These results corroborate our hypothesis that Colombian landraces represent an ancestral form in-between HEMP, from which they descend ((Warf 2014), and MJ, to which they may have made an outsize contribution in the 1960s, 70s, and 80s ((Parsons and Gilmore 1999).

#### pan-NLRome

A total of 7933 predicted R-proteins were grouped into orthogroups with DeepClust and Orthofinder, which identified 96 and 207 orthogroups. We found that 65% or 66% of NLR alleles qualified as CORE; i.e., their containing orthogroups occurred in at least 95% of genotypes (Supp. Tables 7 and 8).

Among the more common alleles, we report fewer clusters, with less SHELL and CLOUD fraction, than two recent reports on the panNLRome in Arabidopsis (Van de Weyer *et al*. 2019; Teasdale *et al*. 2024), both of which deploy more sophisticated, transcriptome-based methods for protein prediction. Similarly to the present study, DIAMOND was used in both for all-by-all alignment; however, the clustering methods varied. Comparing our OrthoFinder results to those of Teasdale et al., our count of 207 OGs was somewhat fewer than the 247 found in Arabidopsis. Van de Weyer et al. used OrthaGogue and found 464 OGs, of which 106, containing 53% of alleles, constituted the CORE, i.e., found in at least 80% of accessions. While still very preliminary, our counting of 65% of alleles as CORE, defined more stringently as OGs present in 95% of accessions, indicates an actual difference in the structure of the two NLRomes. (Using the 80% threshold of Van de Weyer et al., we find that 78% of alleles are CORE.) Consequently, we speculate that a dioecious anemophile such as Cannabis tends to lose orthogroups, either by drift or selection, at a reduced rate relative to an inbreeder such as Arabidopsis.

Other approaches, such as transcriptomics, might reveal unusual alleles not predicted here, and so we do not claim these results to be definitive. PCA and admixture both show strong differentiation between MJ and hemp; yet, at the (predicted) protein level 100% of CORE and SHELL orthogroups include members from both (Supp. Tables 7 and 8). Anecdotally, MJ is broadly considered to be more susceptible than HEMP, as evidenced, and likely exacerbated, by the predominance of protected culture for the former and open-field cultivation for the latter. Our preliminary analysis indicates that HEMP does not contain any unique NLR orthogroups, and so we suggest, tentatively, that its more resistant nature may not be R-gene mediated.

## Conclusions

The study of pangenomes broadens many fields of inquiry that are relevant for both basic and applied research. By evaluating the structural variation that exists across a diverse population, one gains insights into presence-absence variation, copy number variation, and chromosomal architecture that may not be detected by mapping reads to an individual reference.

Here, we show that dual assemblies may be derived from ONT R9 reads without needing parental data, a result not previously reported, but one also already made obsolete with the advent of the R10 pore. Still, our assemblies offer contiguity and gene content comparable to more modern methods, and can help to clarify the arrangement of certain repetitive regions. Assembling dual assemblies from ONT reads is not a solved problem, and perhaps our extensive optimization could inform future efforts.

Our phylogenomic analysis, based on whole genome alignments, is rapid and facile, and avoids the error term associated with mapping reads to repetitive regions. With the continuation of extremely stringent THC limits for hemp production, understanding how ancestry influences its accumulation remains important. Finally, our pan-NLRome sheds some light on how allelic variation may be more likely to persist when inbreeding is suppressed.

For the future, incorporating these 8 new assemblies into comprehensive pangenome graphs should offer new insights into the evolution of this important plant, and permit its continued development as a valuable crop and tractable model system.

## Declarations

We thank Juan Guillermo Torres Hurtado for facilitating the use of the ZINE high performance computing cluster, of the Pontificia Universidad Javeriana.

We gratefully acknowledge the provision of sequence data and permission to publish granted to us by Hawthorne Gardening Company.

## Acknowledgements

We thank David Konkin for preparing and sequencing the genomic libraries. The sequencing was carried out at the National Research Council of Canada’s Aquatic and Crop Resource Development (ACRD) Research Centre in Saskatoon, Saskatchewan.

We thank Lighthouse Genomics and Hawthorne Gardening Company for providing the sequence data and permitting its publication.

We thank Juan Guillermo Torres Hurtado for facilitating the use of the Pontificia Universidad Javeriana’s ZINE high-performance computing cluster.

We dedicate this manuscript to the memory of Anders Gonçalves da Silva.

## Authors’ contributions

AGdS conceived the project and selected the genotypes for sequencing. BP developed the pipeline to assemble, annotate, and compare the genomes, and drafted the manuscript. WT supervised the project and edited the manuscript.

## Supplemental Figures

**Supplemental Figure 1.**
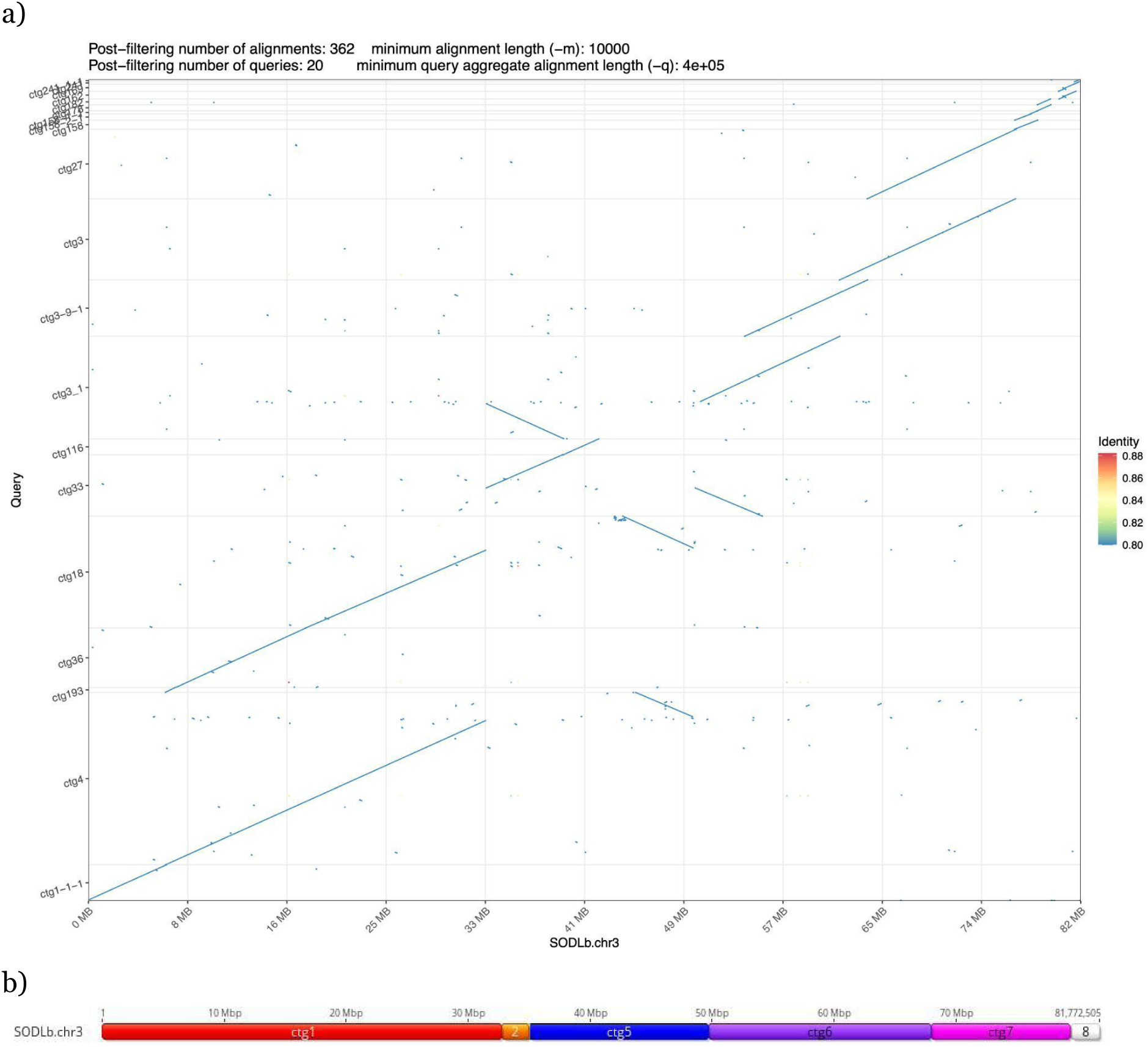
(a) Alignment of a representative pair of diploid contigs against Sour Diesel B chromosome 3, and (b) contig boundaries of the SODLb contigs comprising the chromosome.

**Supplemental Figure 2.**
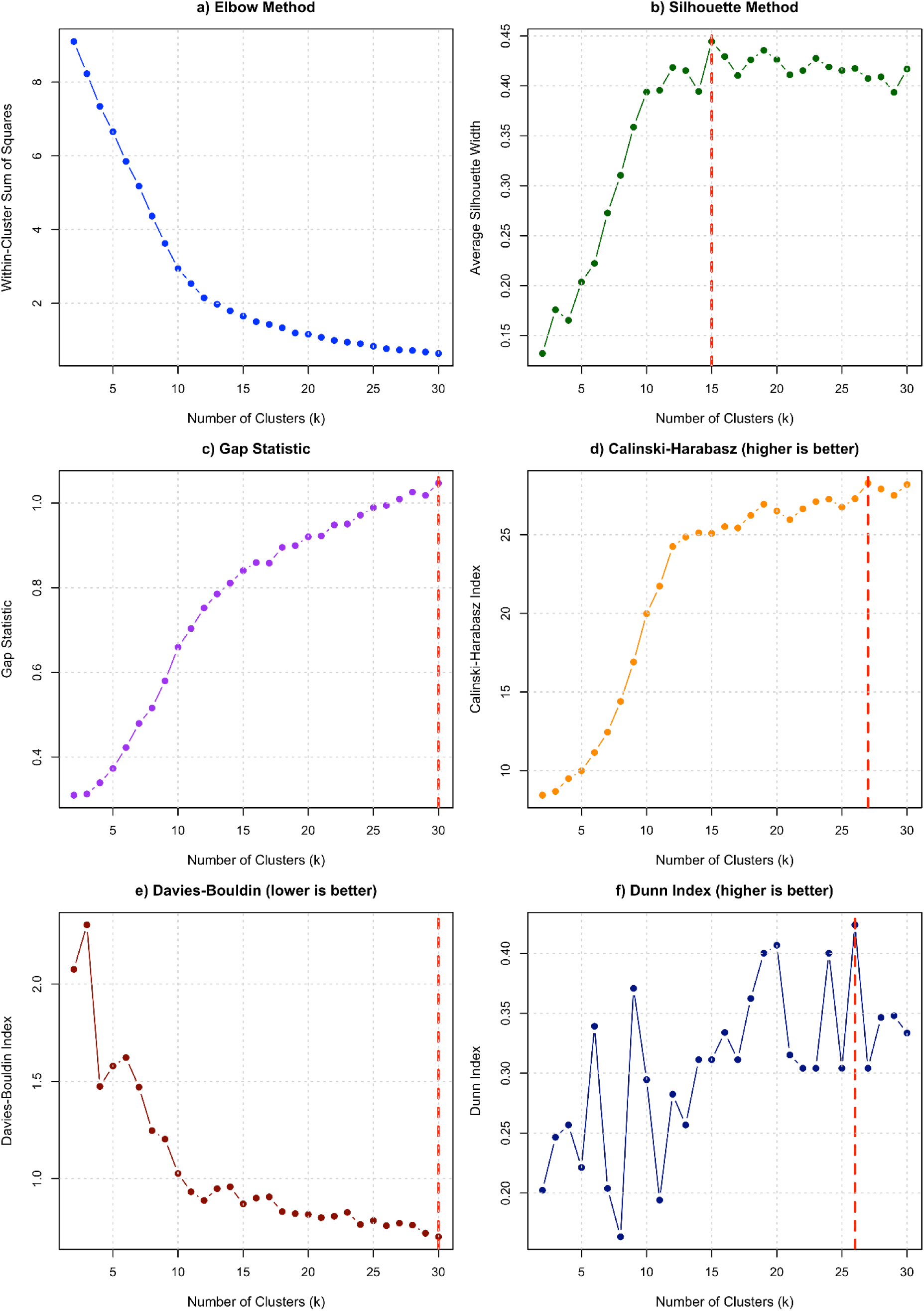
Estimates of subpopulation number among the 85 genotypes used for phylogenomic analysis.

**Supplemental Figure 3.**
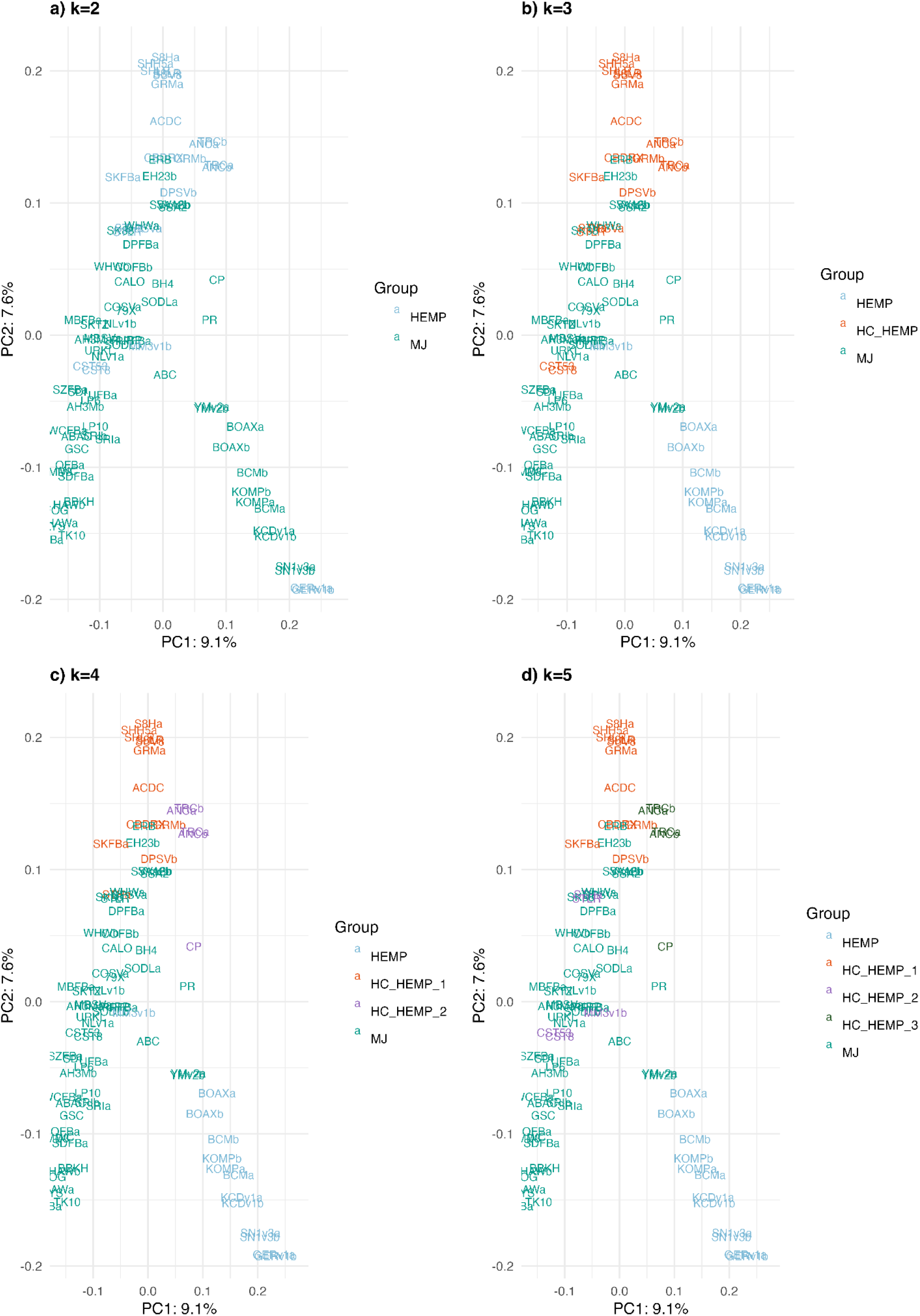
Clustering by Partitioning Around Medoids (PAM), with k from 2 to 5. Note that EH23, SSA, SVA (all HC_HEMP), and YMv2 (HEMP) consistently group with MJ.

**Supplemental Figure 4.**
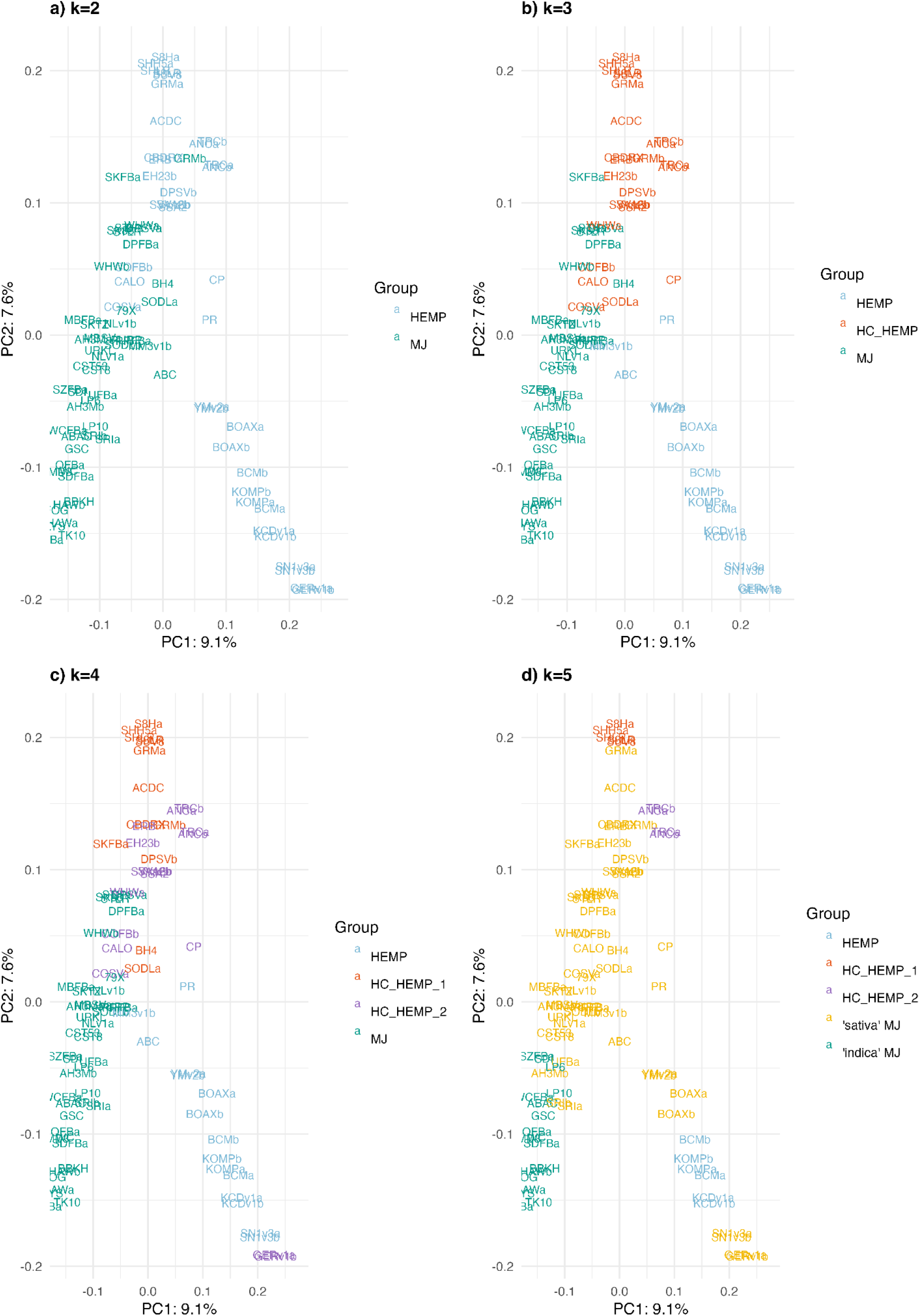
Clustering with Gaussian Mixed Model (GMM) with k from 2 to 5. Note that in no case do HEMP, MJ, and HC_HEMP separate cleanly.

**Supplemental Figure 5.**
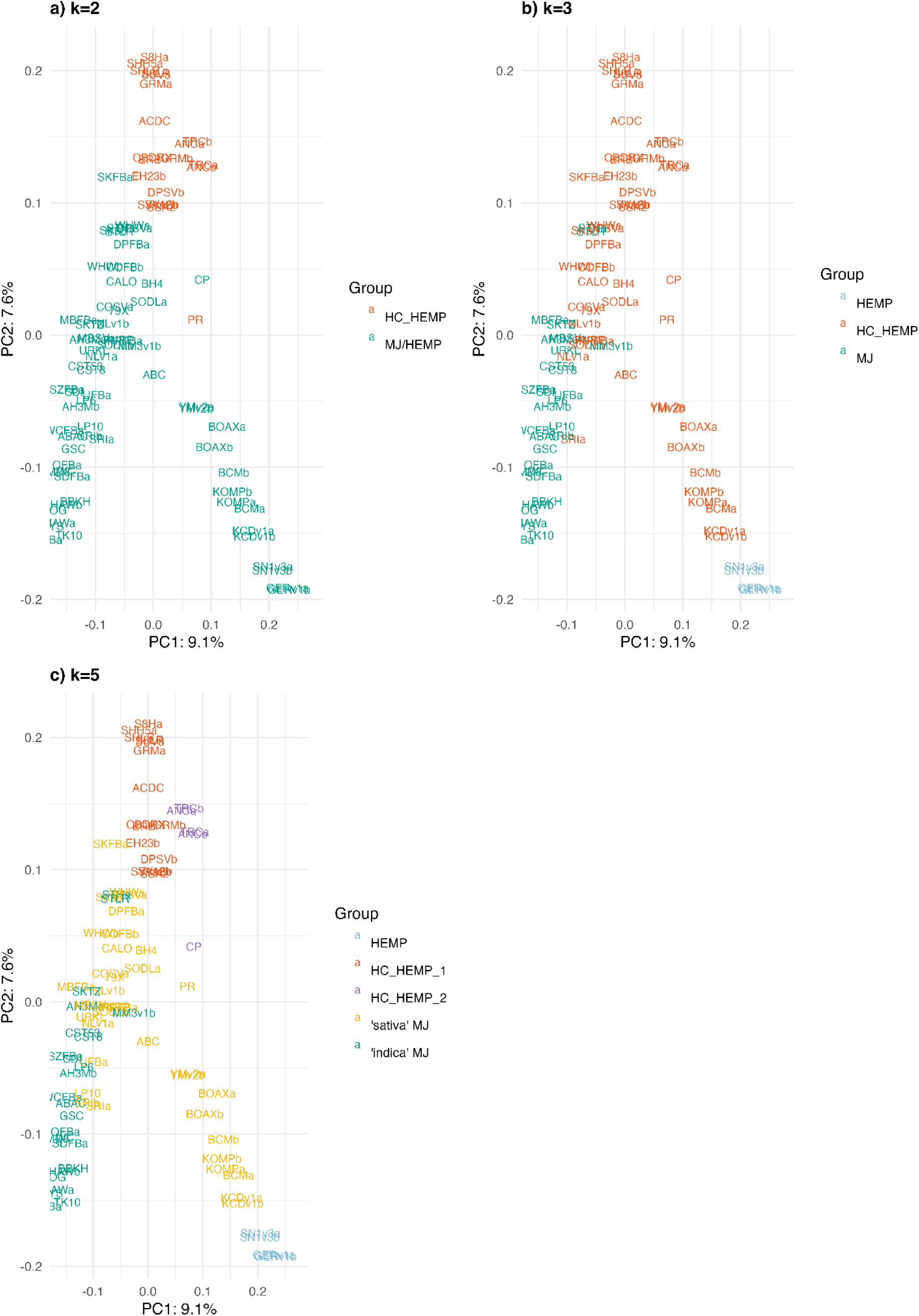
K-means clustering with k of 2, 3, or 5. Note that ‘sativa’ MJ tends to conflate with either HEMP or HC_HEMP.

**Supplemental Figure 6.**
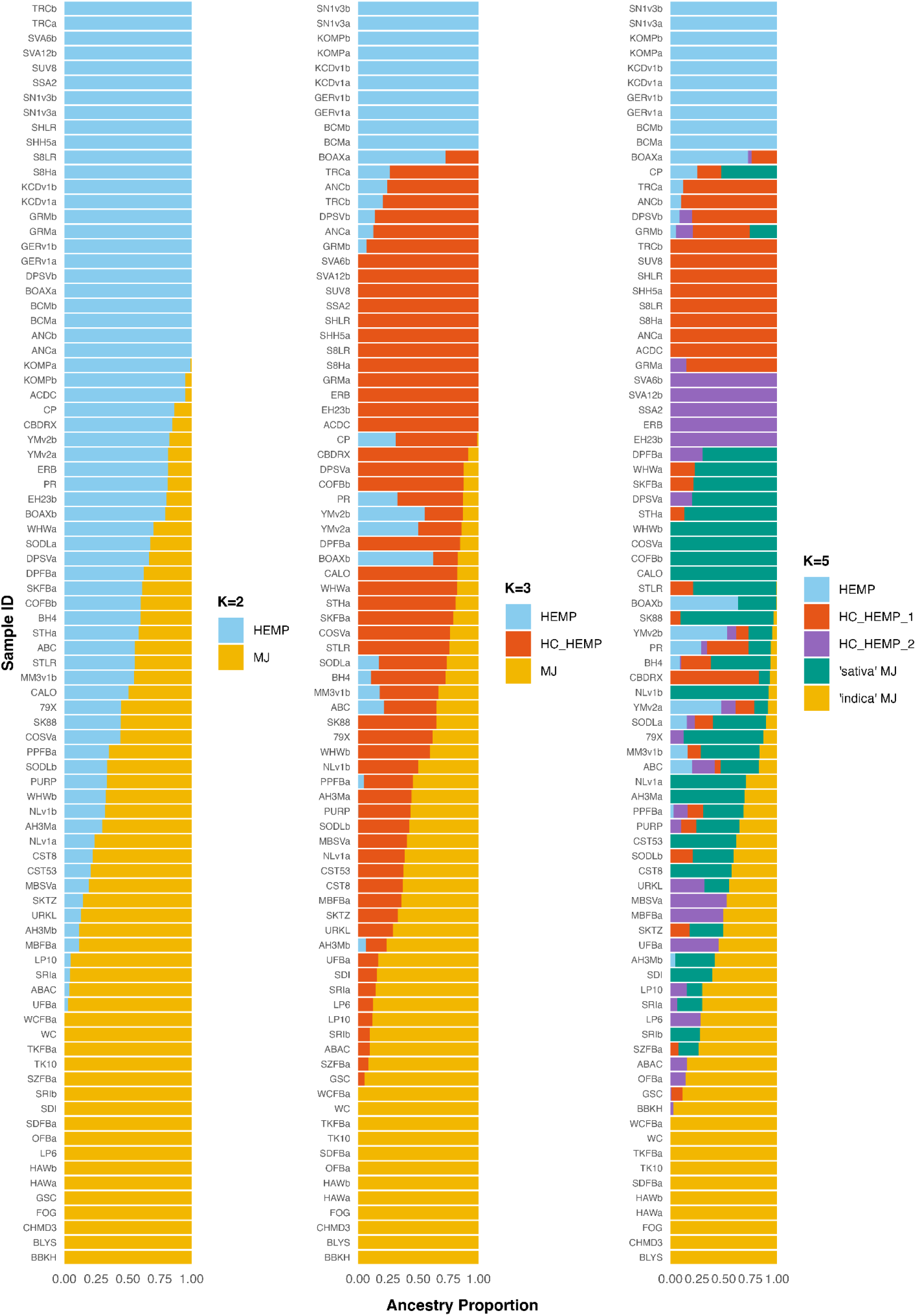
ADMIXTURE plots with k of 2, 3, or 5.

## References

Alexander DH, Lange K. Enhancements to the ADMIXTURE algorithm for individual ancestry estimation. BMC Bioinformatics 2011;12:246.

Booth JK, Page JE, Bohlmann J. Terpene synthases from Cannabis sativa. PLoS One 2017;12:e0173911.

Booth JK, Yuen MMS, Jancsik S et al. Terpene Synthases and Terpene Variation in Cannabis sativa. Plant Physiol 2020;184:130–47.

Braich S, Baillie RC, Spangenberg GC et al. A new and improved genome sequence of Cannabis sativa. Gigabyte 2020;2020:1–13.

Buchfink B, Ashkenazy H, Reuter K et al. Sensitive clustering of protein sequences at tree-of-life scale using DIAMOND DeepClust. bioRxiv 2023:2023.01.24.525373.

Bushnell. BBTools: a suite of fast, multithreaded bioinformatics tools designed for analysis of DNA and RNA sequence data. Joint Genome Institute 2018.

Chen X, Guo H-Y, Zhang Q-Y et al. Whole-genome resequencing of wild and cultivated cannabis reveals the genetic structure and adaptive selection of important traits. BMC Plant Biol 2022;22:371.

Coombe L, Kazemi P, Wong J et al. Multi-genome synteny detection using minimizer graph mappings. bioRxiv 2024:2024.02.07.579356.

Coombe L, Nikolić V, Chu J et al. ntJoin: Fast and lightweight assembly-guided scaffolding using minimizer graphs. Bioinformatics 2020;36:3885–7.

Coombe L, Warren RL, Birol I. ntSynt-viz: Visualizing synteny patterns across multiple genomes. bioRxiv 2025:2025.01.15.633221.

Craen S de, Commandeur JJF, Frank LE et al. Effects of group size and lack of sphericity on the recovery of clusters in K-means cluster analysis. Multivariate Behav Res 2006;41:127–45.

Danecek P, Auton A, Abecasis G et al. The variant call format and VCFtools. Bioinformatics 2011;27:2156–8.

Eddy SR. Profile hidden Markov models. Bioinformatics 1998;14:755–63.

Emms DM, Kelly S. OrthoFinder: phylogenetic orthology inference for comparative genomics. Genome Biol 2019;20:238.

English AC, Menon VK, Gibbs RA et al. Truvari: refined structural variant comparison preserves allelic diversity. Genome Biol 2022;23:271.

Gao S, Wang B, Xie S et al. A high-quality reference genome of wild Cannabis sativa. Hortic Res 2020;7:73.

Ghurye J, Rhie A, Walenz BP et al. Integrating Hi-C links with assembly graphs for chromosome-scale assembly. PLoS Comput Biol 2019;15:e1007273.

Goel M, Schneeberger K. plotsr: visualizing structural similarities and rearrangements between multiple genomes. Bioinformatics 2022;38:2922–6.

Goel M, Sun H, Jiao W-B et al. SyRI: finding genomic rearrangements and local sequence differences from whole-genome assemblies. Genome Biol 2019;20:277.

Grassa CJ, Weiblen GD, Wenger JP et al. A new Cannabis genome assembly associates elevated cannabidiol (CBD) with hemp introgressed into marijuana. New Phytol 2021, DOI: 10.1111/nph.17243.

Hamming RW. Error detecting and error correcting codes. Bell Syst Tech J 1950;29:147–60.

Hennink S. Optimisation of breeding for agronomic traits in fibre hemp (Cannabis sativa L.) by study of parent-offspring relationships. Euphytica 1994;78:69–76.

Hoff KJ, Stanke M. Predicting genes in single genomes with AUGUSTUS. Curr Protoc Bioinformatics 2019;65:e57.

Hou Y, Wang L, Pan W. Comparison of Hi-C-Based Scaffolding Tools on Plant Genomes. Genes 2023;14, DOI: 10.3390/genes14122147.

Jiang H. paf2dotplot: Draw a Dot Plot from a Paf Alignment. Github, 2021.

Jin D, Henry P, Shan J et al. Classification of cannabis strains in the Canadian market with discriminant analysis of principal components using genome-wide single nucleotide polymorphisms. PLoS One 2021;16:e0253387.

Kang X, Xu J, Luo X et al. Hybrid-hybrid correction of errors in long reads with HERO. bioRxiv 2023:2023.11.10.566673.

Kourelis J, Sakai T, Adachi H et al. RefPlantNLR is a comprehensive collection of experimentally validated plant disease resistance proteins from the NLR family. PLoS Biol 2021;19:e3001124.

Kronenberg ZN, Rhie A, Koren S et al. Extended haplotype-phasing of long-read de novo genome assemblies using Hi-C. Nat Commun 2021;12:1935.

Li H. Minimap2: pairwise alignment for nucleotide sequences. Bioinformatics 2018;34:3094–100.

Li H. Minipileup: Simple Pileup-Based Variant Caller. Github, 2024.

Lim JD. Cannabis sativa genome assembly ASM2916894v1. Cannabis sativa Isolate KNU-18-1 (Cultivar: Pink Pepper), Whole Genome Sequencing Project, GenBank 2023.

Lynch RC, Padgitt-Cobb LK, Garfinkel AR et al. Domesticated cannabinoid synthases amid a wild mosaic cannabis pangenome. Nature 2025:1–10.

Lynch RC, Vergara D, Tittes S et al. Genomic and Chemical Diversity in Cannabis. CRC Crit Rev Plant Sci 2016;35:349–63.

McPartland JM. Cannabis sativa and Cannabis indica versus “Sativa” and “Indica.” Cannabis Sativa L. -Botany and Biotechnology. Cham: Springer International Publishing, 2017, 101–21.

McPartland JM, Hegman W, Long T. Cannabis in Asia: its center of origin and early cultivation, based on a synthesis of subfossil pollen and archaeobotanical studies. Veg Hist Archaeobot 2019;28:691–702.

de Meijer EPM, Bagatta M, Carboni A et al. The inheritance of chemical phenotype in Cannabis sativa L. Genetics 2003;163:335–46.

Nie F, Ni P, Huang N et al. De novo diploid genome assembly using long noisy reads. Nat Commun 2024;15:2964.

NORML. Scientists have published over 30,000 papers about cannabis over the past decade. NORML 2023.

Obinu L, Trivedi U, Porceddu A. Benchmarking of Hi-C tools for scaffolding plant genomes obtained from PacBio HiFi and ONT reads. Front Bioinform 2024;4:1462923.

Ouchi S, Kajitani R, Itoh T. GreenHill: a de novo chromosome-level scaffolding and phasing tool using Hi-C. Genome Biol 2023;24:162.

Ou S, Scheben A, Collins T et al. Differences in activity and stability drive transposable element variation in tropical and temperate maize. Genome Res 2024;34:1140–53.

Ou S, Su W, Liao Y et al. Benchmarking transposable element annotation methods for creation of a streamlined, comprehensive pipeline. Genome Biol 2019;20:275.

Parsons JJ, Gilmore RL. Colombia -Drug Trafficking, Guerrilla Warfare, Conflict. Encyclopedia Britannica 1999.

Phylos Bioscience, Inc. Cannabis sativa genome assembly Csat_AbacusV2. Genome assembly Csat_AbacusV2 2022.

Pike B, Kozik A, Terán W. A trio-binning approach for genome assembly reveals extensive structural variation between two Cannabis cultivars: Punto Rojo and Cherry Pie. G3 Genes|Genomes|Genetics 2025, DOI: 10.1093/g3journal/jkaf286.

Porubsky D, Ebert P, Audano PA et al. Fully phased human genome assembly without parental data using single-cell strand sequencing and long reads. Nat Biotechnol 2021;39:302–8.

Ramsay L, Koh CS, Kagale S et al. Genomic rearrangements have consequences for introgression breeding as revealed by genome assemblies of wild and cultivated lentil species. Plant Biology 2021.

Ren G, Zhang X, Li Y et al. Large-scale whole-genome resequencing unravels the domestication history of Cannabis sativa. Sci Adv 2021;7, DOI: 10.1126/sciadv.abg2286.

Roach MJ, Schmidt SA, Borneman AR. Purge Haplotigs: allelic contig reassignment for third-gen diploid genome assemblies. BMC Bioinformatics 2018;19:1–10.

Short DJ. Blueberry F5. DJ Short Seeds 2019.

Shumate A, Salzberg SL. Liftoff: accurate mapping of gene annotations. Bioinformatics 2021;37:1639–43.

Steuernagel B, Witek K, Krattinger SG et al. The NLR-Annotator tool enables annotation of the intracellular immune receptor repertoire. Plant Physiol 2020, DOI: 10.1104/pp.19.01273.

Sur A, Myler PJ. A benchmark of Hi-C scaffolders using reference genomes and de novo assemblies. bioRxiv 2022:2022.04.20.488415.

Teasdale LC, Murray KD, Collenberg M et al. Pangenomic context reveals the extent of intraspecific plant NLR evolution. bioRxiv 2024:2024.09.02.610789.

Van de Weyer A-L, Monteiro F, Furzer OJ et al. A Species-Wide Inventory of NLR Genes and Alleles in Arabidopsis thaliana. Cell 2019;178:1260–72.e14.

Vaser R, Sović I, Nagarajan N et al. Fast and accurate de novo genome assembly from long uncorrected reads. Genome Res 2017;27:737–46.

Vasimuddin M, Misra S, Li H et al. Efficient architecture-aware acceleration of BWA-MEM for multicore systems. 2019 IEEE International Parallel and Distributed Processing Symposium (IPDPS). IEEE, 2019, 314–24.

Wang S, Zhong X, Cheng Y et al. Pan-genome analysis of cannabis sativa: Insights on genomic diversity, evolution, and environment adaption. Int J Mol Sci 2025;26:8354.

Warf B. High Points: An Historical Geography of Cannabis. Geogr Rev 2014;104:414–38.

Warren RL, Coombe L, Mohamadi H et al. ntEdit: scalable genome sequence polishing. Bioinformatics 2019;35:4430–2.

Wickham H. Ggplot2: Ggplot2. Wiley Interdiscip Rev Comput Stat 2011;3:180–5.

Wick R. Porechop: Adapter Trimmer for Oxford Nanopore Reads. Github, 2018.

Woods P, Campbell BJ, Nicodemus TJ et al. Quantitative Trait Loci Controlling Agronomic and Biochemical Traits in Cannabis sativa. Genetics 2021, DOI: 10.1093/genetics/iyab099.

Zhang Q, Chen X, Guo H et al. Latitudinal adaptation and genetic insights into the origins of Cannabis sativa L. Front Plant Sci 2018;9:1876.

Zheng Z, Li S, Su J et al. Symphonizing pileup and full-alignment for deep learning-based long-read variant calling. Nature Computational Science 2022:2021.12.29.474431.

Zhou C, McCarthy SA, Durbin R. YaHS: yet another Hi-C scaffolding tool. Bioinformatics 2023;39, DOI: 10.1093/bioinformatics/btac808.

Zimmermann L, Stephens A, Nam S-Z et al. A completely reimplemented MPI Bioinformatics Toolkit with a new HHpred server at its core. J Mol Biol 2018;430:2237–43.

